# Adaptive resetting of tuberoinfundibular dopamine (TIDA) network activity during lactation in mice

**DOI:** 10.1101/797159

**Authors:** Carolina Thörn Pérez, Jimena Ferraris, Josina Anna van Lunteren, Arash Hellysaz, María Jesús Iglesias, Christian Broberger

## Abstract

Giving birth triggers a wide repertoire of physiological and behavioural changes in the mother to enable her to feed and care for her offspring. These changes require coordination and are often orchestrated from the central nervous system, through as of yet poorly understood mechanisms. A neuronal population with a central role in puerperal changes is the tuberoinfundibular dopamine (TIDA) neurons that control release of the pituitary hormone, prolactin, which triggers key maternal adaptations, including lactation and maternal care. Here, we used Ca^2+^ imaging on mice from both sexes and whole-cell recordings on female mouse TIDA neurons *in vitro* to examine if they adapt their cellular and network activity according to reproductive state. In the high-prolactin state of lactation, TIDA neurons shift to faster membrane potential oscillations, a reconfiguration that reverses upon weaning. During the estrous cycle, however, which includes a brief, but pronounced, prolactin peak, oscillation frequency remains stable. An increase in the hyperpolarization-activated mixed cation current, I_h_, possibly through unmasking as dopamine release drops during nursing, may explain the reconfiguration of TIDA rhythms. These findings identify a reversible plasticity in hypothalamic network activity that can serve to adapt the dam for motherhood.

**Significance Statement:** Motherhood requires profound behavioural and physiological adaptations to enable caring for offspring, but the underlying CNS changes are poorly understood. Here, we show that during lactation, neuroendocrine dopamine neurons, the “TIDA” cells that control prolactin secretion, reorganize their trademark oscillations to discharge in faster frequencies. Unlike previous studies, which typically have focused on structural and transcriptional changes during pregnancy and lactation, we demonstrate a functional switch in activity and one that, distinct from previously described puerperal modifications, reverses fully upon weaning. We further provide evidence that a specific conductance – I_h_ - may underlie the altered network rhythm. These findings identify a new facet of maternal brain plasticity at the level of membrane properties and consequent ensemble activity.

## INTRODUCTION

The mother who has just given birth undergoes a series of physiological changes. Parallel to recalibration after the extensive pregnancy-associated organ modifications, physical and behavioural adaptive modifications enable caring for the offspring (Brunton and Russell, 2008). These changes are orchestrated by an altered blood profile of hormones, which, in turn, likely require reconfiguration at the master level of the endocrine system, the hypothalamus-pituitary axes. There is abundant evidence that the transcriptional repertoire of neuroendocrine populations undergoes dramatic revision during the bearing and rearing of children (see (Augustine et al., 2018; Grattan et al., 2008). Far less is, however, known about if, and how, network properties of the hypothalamic neurons that control the pituitary are subject to adaptive resetting during nursing.

Among the hormones that trigger and sustain the maternal state, prolactin plays a central role (Freeman et al., 2000). Originally named for its ability to trigger crop milk production in pigeons (Stricker, 1928), prolactin is now understood to induce a wide range of the *post-partum* physiological changes in the mother (see (Grattan et al., 2008). These include modifications of ovulatory function (Cohen-Becker, 1986; Fox, 1987), libido (Dudley et al., 1982), maternal behaviour (Bridges, 1985; Brown et al., 2017)), the immune system (see (Dorshkind and Horseman, 2000)), and metabolism and feeding (Sauve, 1996).

Blood prolactin levels exhibit a marked dynamic profile, driven primarily by reproductive state (Ronnekleiv and Kelly, 1988). While lactation corresponds to a sustained hyperprolactinemic state, beginning during late pregnancy in rodents (Sairenji et al., 2017), circulating prolactin is low under baseline conditions in females and, even more so, in males (Guillou et al., 2015; Kalyani et al., 2016). A brief, albeit pronounced, peak appears within the estrous cycle in the transition from proestrus to estrus (Butcher et al., 1974), enabling the maintenance of the corpus luteum and successful implantation (Morishige and Rothchild, 1974).

The dominant control of pituitary prolactin secretion is exerted by neuroendocrine tuberoinfundibular dopamine (TIDA) neurons located in the dorsomedial hypothalamic arcuate nucleus (dmArc; see (Lyons and Broberger, 2014)). These neurons powerfully inhibit prolactin exocytosis by releasing dopamine via the portal capillary vessels to act on dopamine D2 receptors (D2Rs) on the lactotroph cells. Dopamine release usually mirrors circulating prolactin to mediate feedback inhibition (Brown et al., 2016; Lerant and Freeman, 1998), but during lactation, this control is proposed to be overridden through downregulated dopamine biosynthesis (Wang et al., 1993), enabling prolactin to remain elevated for long periods to enable nursing (see (Grattan, 2015; Le Tissier et al., 2015; Yip et al., 2019)).

Yet, neurons calibrate their action not merely by the number of receptors they express and transmitter they produce, but equally important by their cellular and network activity patterns. The issue of adaptive control of TIDA neurons is interesting, not only for their role in reproduction and nursing, but also for the intriguing pattern of their discharge activity, organized as slow, regular membrane potential oscillations (Lyons et al., 2010). These oscillations can be observed by whole-cell patch clamp recording as well as Ca^2+^ imaging in both rats and mice, albeit with species-specific features in network configuration (Stagkourakis et al., 2018). TIDA network activity can be modulated by several hormones and transmitters implicated in prolactin control (Briffaud et al., 2015; Lyons et al., 2016; Stagkourakis et al., 2016). In the pituitary, the ensemble activity of lactotrophs has been shown to be reorganized as a female enters nursing, with remarkably long-lasting effects (Hodson et al., 2012; Zhang and van den Pol, 2015). If - and how - the network activity of TIDA neurons, the master control of the lactotropic axis, is reconfigured as a consequence of reproductive state remains, however, largely unknown. Here, we used Ca^2+^ imaging and whole-cell recordings, combined with *in situ* hybridization, in mice to determine the cellular and network activity of these neurons during different reproductive states. We demonstrate a reversible shift in ensemble activity during lactation, and identify a candidate underlying mechanism in altered membrane properties.

## MATERIAL AND METHODS

### ANIMALS

Experiments were approved by the local animal research ethical committee and *Stockholms Norra Djurförsöksetiska Nämnd*.

Experiments were performed in F1 adult (4-6 months) female and male transgenic mice (on a C57BL/6J background) expressing GCaMP3 or tdTomato under the control of the dopamine transporter (DAT; (Ekstrand et al., 2007). Females were divided in different groups according to the estrogen cycle: proestrus, estrus, diestrus/metestrus (determined by the cytology of the vaginal smears; (McLean et al., 2012). For the lactating studies, dams were used between days 7 and 10 of lactation. For weaning studies, pups were removed from the dams at postnatal day 21, which were used for recordings three weeks later.

Animals were anesthetized with pentobarbital, decapitated and the brain was dissected and sliced at 4°C in slicing solution of the following composition (in mM): sucrose (213), KCl (2), NaH_2_PO_4_ (1.3), NaHCO_3_ (26), CaCl_2_ (2), MgCl_2_ (2) and glucose (10). Two hundred and fifty µm thick coronal hypothalamic slices were cut on a vibratome. Slices where continuously perfused with oxygenated artificial cerebrospinal fluid (aCSF) containing the following (in mM): NaCl (127), KCl (2), NaH_2_PO_4_ (1.3), NaHCO_3_ (26), CaCl_2_, (2.4), MgCl_2_ (1.3) and glucose (10). Three to five slices per animal –with a minimum of four oscillating cells per slice-were used for the network-mapping experiments.

### IMMUNOFLUORESCENCE

Adult male mice (n=4) were anesthetized and perfused via the ascending aorta with a fixative containing 4% paraformaldehyde and 0.2% picric acid. Brains were dissected, cryo-protected in sucrose, rapidly frozen with CO_2_ gas and sectioned on a cryostat at 14 μm thickness. Tissue sections spanning the entire rostro-caudal length of the Arc were processed for immunofluorescence (Foo et al., 2014), using primary anti-tyrosine hydroxylase (TH) antiserum (1:2,000; raised in rabbit; AB152; Millipore, RRID:SCR_008983) combined with anti-green fluorescence protein (GFP) antiserum (1:4000; raised in chicken; GFP-1020; Aves Labs Cat# GFP-1020, RRID:AB_10000240). TH was visualized with Alexa-594 conjugated secondary antisera (1:500; raised in donkey; A21207; Thermo Fisher Scientific) and GFP was visualized with tyramide signal amplification (TSA) Plus (Perkin Elmer) using horseradish peroxidase (HRP)- conjugated secondary antisera (1:500; raised in goat; 103-035-155; Jackson ImmunoResearch) and fluorescein tyramide substrate. Tissue sections were mounted with 2.5% 1,4-Diazabicyclo[2.2.2]octane (DABCO) in 100% glycerol. Fluorescence micrograph montages were automatically generated by taking consecutive pictures with a Zeiss Axio Imager M1 and merging them using MBF Neurolucida computer software. Confocal micrographs were acquired with an Olympus FV1000 microscope and visualized in BitPlane® Imaris. For final images, brightness, contrast and sharpness were adjusted digitally.

### RNASCOPE ASSAY

Adult mice (lactating and females in estrous) were deeply anesthetized with isoflurane. The brain was removed and rapidly frozen on dry ice. Brains were stored at −80°C until sectioning. Coronal sections (14-20 µm) of the hypothalamus were cut on a cryostat (CryoStar NX70, Thermo Scientific) and mounted on SuperFrost Plus (Fisher Scientific) slides. Slides were processed for fluorescence *in situ* hybridization of multiple target RNAs simultaneously according to manufacturer protocols for RNAScope (Advanced Cell Diagnostics). Briefly, sections were post fixed in 10% neutral buffered formalin, washed and dehydrated in sequential concentrations of ethanol (50, 70 and 100%) at room temperature. Samples were treated with protease, then incubated for two hours at 40°C in the presence of target probes to allow for hybridization. A series of three amplification steps is necessary to provide substrate for target fluorophore labeling. After labeling, samples were counterstained. Prolong Gold Antifade Mountant (Invitrogen) was applied to each slide prior to cover slipping.

*Imaging.* Images were acquired by confocal laser scanning microscope (LSM710 META, Zeiss). High-resolution z-stack confocal images were taken at 0.3 μm intervals.

### PROLACTIN LEVEL ANALYSIS

Mice were anesthetized with an i.p. dose of pentobarbital and trunk blood samples were collected after decapitation. The serum was obtained by leaving the blood to clot at room temperature for 30 min and centrifugation at 4°C for 15 min (10.000 rpm). The supernatant was aliquoted and stored at −80°C. Prolactin serum levels were measured by bead-based sandwich immunoassay (Qundos et al., 2014). Affinity reagents and the mouse recombinant prolactin protein, purchased to RnD Biosystems Douset kit (RnD DY1445), were used to quantify prolactin levels in serum. In brief, a goat anti-mouse prolactin capture antibody, as well as goat IgG, were covalently coupled to colour-coded magnetic beads (MagPlex® microspheres), which, together with plain beads (as negative control), composed the bead-based array. The coupling efficiency for the antibody was determined via R-phycoerythrin-labeled anti-goat IgG antibody (Jackson ImmunoResearch).

Serum samples were diluted 1:5 in assay buffer and then 45 μl were combined with 5 μl of the bead array in microtiter plates (Greiner). Samples incubated for 2 hours on a shaker at RT and 650 rpm. Beads were washed on a magnet with PBS-T using a plate washer (EL406, BioTek). This was followed by 1 h incubation of 70 ng/ml biotinylated detection antibody (part B-DY1445). Beads were washed again, and 0.5 μg/ml R-phycoerythrin-labeled streptavidin (Invitrogen) in PBS-T was added and incubated for 30 min. Finally, after washes samples were measured in PBS-T using the MAGPIX^TM^ instrument (Luminex Corp.) Median fluorescence Intensity (MFI) and a standard curve were obtained for all samples. A 5PL parameter logistic curve was generated to interpolate MFI into protein concentration (ng/ml).

### CALCIUM IMAGING

The activity of oscillating neurons was recorded in a chamber that was continuously perfused with oxygenated aCSF and maintained at approx. 36° C, on a Zeiss microscope system. Fluorescence images were captured with a CCD camera (Evolve, Photometrics) monitored by fluorescence changes under imaging acquisition in Image J (NIH) or Metafluor software. Imaging frames of 512×512 pixels were acquired at 4 Hz, during 90 seconds using a 20× water-immersion objective (Zeiss), and image sequences were analysed with custom software written in ImageJ (National Center for Microscopy and Imaging Research: ImageJ Mosaic Plug-ins, RRID:SCR_001935) and MATLAB (Math Works, MATLABAB, RRID:SCR_001622).

Individual cells were marked as regions of interest (ROIs) from the image sequences *i.e,* all positive cells in one plane. Consistent with previous studies in mice, both oscillating and non-oscillating DAT-GCaMP3 cells were found (Stagkourakis et al., 2018; Zhang and van den Pol, 2015). Our focus in this study was on the oscillating population, which was selected by the fold difference in the intensity signal (≥ 1%) and the rhythmicity index (defined below; ≥ 0.2).

The recorded intensity signals were presented as relative changes of fluorescence in each of the selected ROIs and expressed as (ΔF/F0), where F0 is the minimum fluorescence value of the ROI ΔF = (F – F0) * 100. The frequency was determined using a power spectrum. The signal was high pass filtered using a cut-off at 0.1 Hz and the rhythmicity index (RI) was defined as RI = β/α, where α is the amplitude of the peak at zero lag and β is the amplitude of the peak at the period of the Ca^2+^ oscillations, usually the second peak in the auto-correlogram (see Figure 3B1) (Levine et al., 2002). Thus, the closer RI is to 1, the more rhythmic is the Ca^2+^ signal. A MATLAB script described in (Smedler et al., 2014) was used to illustrate and determine the correlation matrix value and correlated cell pairs were defined as the ones with autocorrelation coefficient ≥ 0.8.

Simultaneous Ca^2+^ imaging and whole-cell patch clamp recording (see below) were performed using a 64× water-immersion objective. Whole-cell recordings were obtained through visualizing DAT-GCaMP3 cells and then adapting them to the differential contrast (DIC) images. For temporal synchronization of acquired images and recorded membrane potential oscillations, camera exposure data indicating the occurrence of each frame was obtained from the imaging system and recorded simultaneously with the intracellular signal using pClamp software (Axon Instruments Inc.).

### ELECTROPHYSIOLOGY

For the whole-cell recording experiments, identical conditions of perfusion rate and temperature were kept as with Ca^2+^ imaging. TIDA neurons – visualized as DAT-GCaMP3 or DAT-tdTomato - were recorded using patch electrodes pulled from borosilicate glass microcapillaries. Whole-cell recordings were performed in current- or voltage-clamp mode using a Multiclamp 700B amplifier (Molecular Devices). Bridge balance and pipette capacitance compensation were automatically adjusted. Liquid junction potential was 15.2 mV and not compensated. Patch pipettes had resistances of 4-7 MΩ and contained the following (in mM): K-gluconate (140), KCl (10), KOH (1), Na_2_ATP (2), HEPES (10). The pH was adjusted to 7.4 with 1 M KOH and the osmolarity to 280 mOsm. Solutions of pharmacological agents were bath-applied at a perfusion rate of 1 ml/min.

Consecutive hyperpolarizing voltage steps (1s, - 140mV to −70 mV) were performed under Na^+^ channel blockade by TTX (500nM; Sigma –Aldrich) in the presence or absence of ZD7288 (Tocris). Drugs were kept as concentrated stocks dissolved in water and kept frozen at −20°C. During the experiments, the drugs were diluted to the final concentration in aCSF and bath applied through the perfusion system.

The signal was amplified, and low pass filtered on-line at 10 kHz and digitized. Data were acquired with Clampex software and analyzed using Clampfit (pCLAMP 10, Molecular Devices), Spike2 4.16 (Cambridge Electronic Design) or WinEDR, Electrophysiological data recorder V3.8.7 (University of Strathclyde) software.

### STATISTICAL ANALYSIS

Summary statistics and the values shown in the figures are reported as standard error of the mean (± SEM) and “n” represents the number of animals, network maps or cells as specified. The Normal distribution of the data was tested using the Kolmogorov-Smirnov test. The significance was determined using ANOVA or t-test with a 95% confidence interval.

## RESULTS

### GCAMP3 DYNAMICS

To access TIDA cells in acute hypothalamic slices, we used a transgenic mouse line expressing the genetically encoded Ca^2+^ indicator, GCaMP3 (under the control of the DAT promoter), which is expressed in dmArc dopamine neurons, *i.e.* the TIDA population (Lorang et al., 1994). GCaMP3 expression was verified by GFP-like immunoreactivity and co-localization of the rate-limiting enzyme in dopamine biosynthesis, tyrosine hydroxylase (TH) (Figures 1A-B).

**FIGURE 1.**
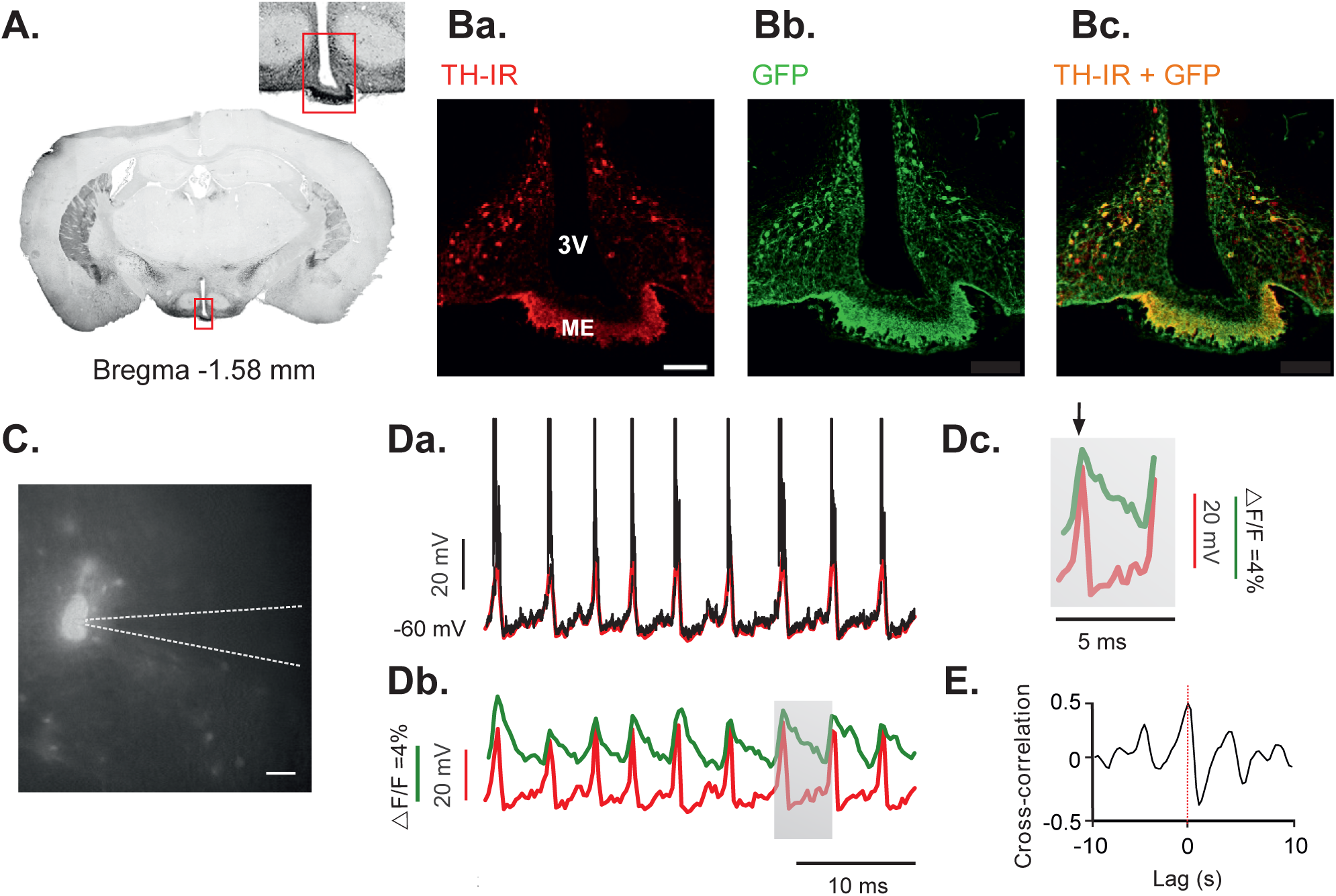
GCAMP3 CA^2+^ IMAGING FAITHFULLY CORRELATES TO ELECTROPHYSIOLOGICALLY RECORDED MEMBRANE POTENTIAL OSCILLATIONS IN TIDA NEURONS. **A.** Fluorescence coronal section of a male DAT–GCaMP3 mouse at the level of the arcuate nucleus. Converted to grayscale and inverted for better contrast. Inset shows higher magnification of marked region. **Ba-c.** Confocal micrographs of marked region in (A) immunostained for tyrosine hydroxylase (TH; red; *Ba*), green fluorescent protein (GFP; green; *Bb*); and merged (*Bc).* (colocalization of signals seen in yellow). **C.** Photomicrograph of a fluorescent DAT-GCaMP3 neuron in the dmArc during whole-cell patch clamp recording. Recording pipette outlined by dashed line. **Da.** Whole-cell current clamp recording from a DAT-GCaMP3 neuron. Down-sampled trace overlaid in red. **Db.** Down-sampled trace from *Da* aligned with the simultaneously recorded Ca^2+^ imaging activity (green). Note coincident activity peaks in both signals. **Dc.** Grey box in *Db* expanded to illustrate high peak-to-peak correlation (black arrow) and the slow decay kinetics of the Ca^2+^ signal compared to the membrane depolarization. **E.** Cross-correlogram between the down sampled membrane potential and the Ca^2+^ activity of trace (*Da*). Scale bar in *Ba*, 100 µm (for *Ba-c*); in *C*, 25 µm. 3V, third ventricle; ME, median eminence.

Voltage-gated Ca^2+^channels mediate a critical step in stimulus-secretion coupling, converting action potential- triggered depolarization to vesicular release of neurotransmitters (Katz and Miledi, 1970). To determine the relationship between GCaMP3-visualized Ca^2+^ fluctuations and membrane potential oscillations in TIDA neurons, we performed simultaneous Ca^2+^ imaging and whole-cell recordings in DAT-GCaMP3 neurons (Figure 1C), revealing a strong correlation with GCaMP3 waveform peaks sharply phase-locked to the TIDA UP states (Figure 1D, n=13/13 cells). The characteristic fast rise and slow decay of the Ca^2+^ signal, is illustrated in Figure 1Db-c.

Cross-correlation analysis of the two signals (Figure 1E) highlights the positive correlation (<0.5, n=13/13) between signal peaks, and the negative correlation between Ca^2+^ signal decay and the slow depolarization phase of the TIDA oscillation. Thus, Ca^2+^ imaging can be used as a reliable readout of the TIDA oscillation in individual neurons as first identified by electrophysiology (Lyons et al., 2010).

### TIDA OSCILLATION FREQUENCY INCREASES DURING LACTATION

To determine if different activity states in the lactotropic axis are paralleled by changes in TIDA network behavior, Ca^2+^ imaging was performed on dmArc slices from DAT-GCaMP3 mice where circulating prolactin is low (juvenile males and females during diestrus/metestrus), transiently elevated (proestrus, estrus) or sustained elevated (lactation). Estrous cycle stages were determined by vaginal smears. The cytology of diestrus and metestrus smears could not consistently be reliably differentiated from each other, but as they are distinct from the other states by the presence of leukocytes, these two states where pooled into one group (D/M). While the characteristic cytological profiles of proestrus and estrus are distinct, there was no significant difference in the oscillation frequency between these states (Figure 2B; p=0.14, unpaired t-test), so data from these two groups were also pooled (P+E). In all physiological conditions the proportion of oscillating cells remained similar (D/M: 53.2±8.6%; P+E 43.7±5.1%; lactating 46.9±9.5%; of all fluorescent cells) and no significant differences between groups; cumulative total average for all groups: 47.9 ± 10.0 %). Figure 2Aa-d show traces of simultaneous Ca^2+^ fluctuations from five cells from a single slice in the experimental groups. The series of images from all groups (males, D/M, estrus and lactating) revealed a large distribution and overlapping dispersal of oscillation frequencies.

**FIGURE 2.**
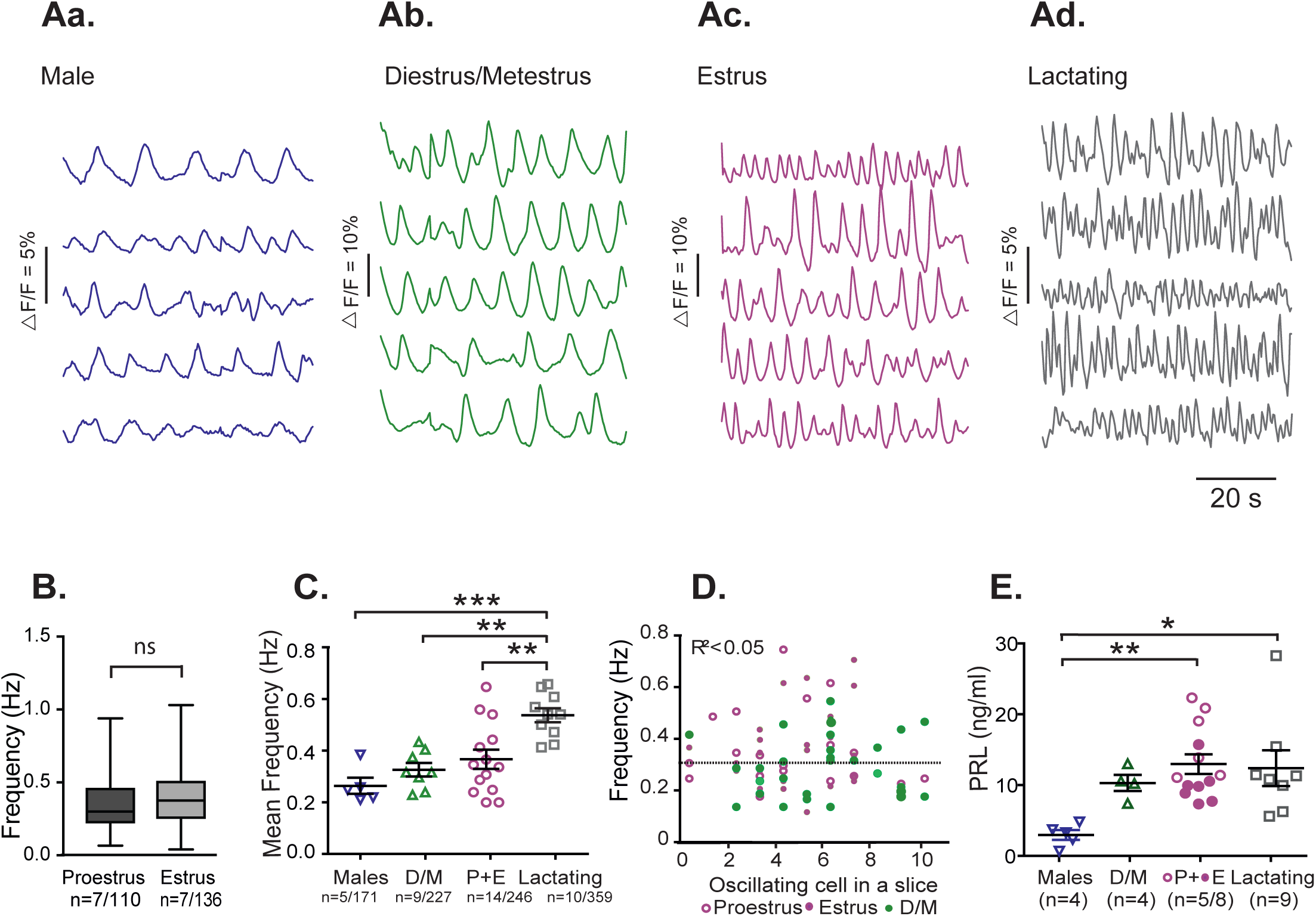
THE FREQUENCY OF TIDA NEURON CA^2+^ OSCILLATIONS INCREASES DURING LACTATION. A. Representative concomitant traces of Ca^2+^fluctuations in DAT-GCaMP3 neurons in the dorsomedial arcuate nucleus (dmArc) from the same slice in adult males (Aa), female in diestrus/metestrus (D/M; Ab), female in estrus (from the proestrus+estrus group; P+E; Ac), and lactating dam (Ad). B. Box plot illustrating the frequency of the Ca^2+^ oscillations in DAT-CGaMP3 neurons in the dmArc recorded from female mice in proestrus and estrus. There is no significant difference (ns) between the two groups (p=0.14). “n” given as the number of animals, the number of cells. C. The mean frequency of oscillating DAT-GCaMP3 neurons in the dmArc in an animal (*i.e.* all oscillating cells, from all slices of a single mouse). The mean frequency was significantly different between the lactating group and the male, D/M and P+E groups. *** P< 0.001, ** P>0.01, all other comparisons between groups are non-significant. “n” given as the number of animals / number of cells, one-way ANOVA with Tukey’s multiple comparison as *post-hoc* test. D. The number of oscillating cells in a slice plotted against the mean oscillation frequency of the same slice. No correlation (Pearson correlation: R^2^ < 0.05) is found, suggesting that the number of active cells within a preparation does not influence the speed of the activity. E. Serum prolactin concentration measured by immunoassay in the experimental groups shown in *B*. * P>0.05, ** P>0.01, all other comparisons between groups are non-significant.

Oscillation frequency was not significantly different in comparisons between adult males (0.25 ± 0.06 Hz) and the estrous cycle groups (D/M = 0.32 ± 0.06 Hz; P+E = 0.36 ± 0.06 Hz). During lactation, TIDA neuron oscillation activity persisted, consistent with a previous report (Romano et al., 2013). The mean frequency of the Ca^2+^ signal fluctuation was, however, significantly faster compared to the other groups (0.53 ± 0.04 Hz; Figure 2C). We found no correlation between the number of oscillating cells in a slice and the mean oscillating frequencies within the nulliparous female population (Figure 2D) suggesting that oscillation period does not depend on active population size.

As summarized above, serum prolactin levels are shaped by TIDA neurons, which exert tonic inhibition on pituitary lactotrophs (see (Grattan, 2015). We performed a bead-based sandwich immunoassay to directly study serum prolactin levels in the mice used in the recordings. Prolactin levels were significantly different between males and the P+E group and between males and the lactating group (Figure 2E).

Taken together, these results suggest that while prolactin levels vary dynamically both during the estrous cycle and during nursing, it is only in conjunction with long-term changes in prolactin output (*i.e.* nursing), and not for more transient elevations (*i.e.* proestrus/estrus), that plastic changes that shift TIDA oscillation frequency appear.

### TIDA RHYTHMICITY DECREASES DURING LACTATION

To further investigate the extent of TIDA network rearrangement across high- and low prolactin states, we conducted a cross-correlation analysis of the Ca^2+^ fluctuations between all oscillating cells in a single network (see *Methods*). Figure 3A shows representative network maps from males, estrous stages and lactation. The colour represents level of cell to cell correlation according to the Ca^2+^ activity profiles. While correlated cell pairs could be observed in all stages, they were rare (one or two per slice), suggesting an overall absence of synchronization. Figure 3C summarizes the matrix value of each map in different physiological states. The differences in functional connectivity were not significant between groups (ANOVA; n corresponds to the number of network maps), suggesting that direct communication between oscillating cells is low (and that a common organizing input is unlikely, at least within the anatomical confines of the slice preparation), and that this condition remains unchanged across the physiological states that we examined. Thus, the state-dependent changes that we observed in oscillation frequency are unlikely to be a consequence of modified connectivity in the network.

**FIGURE 3.**
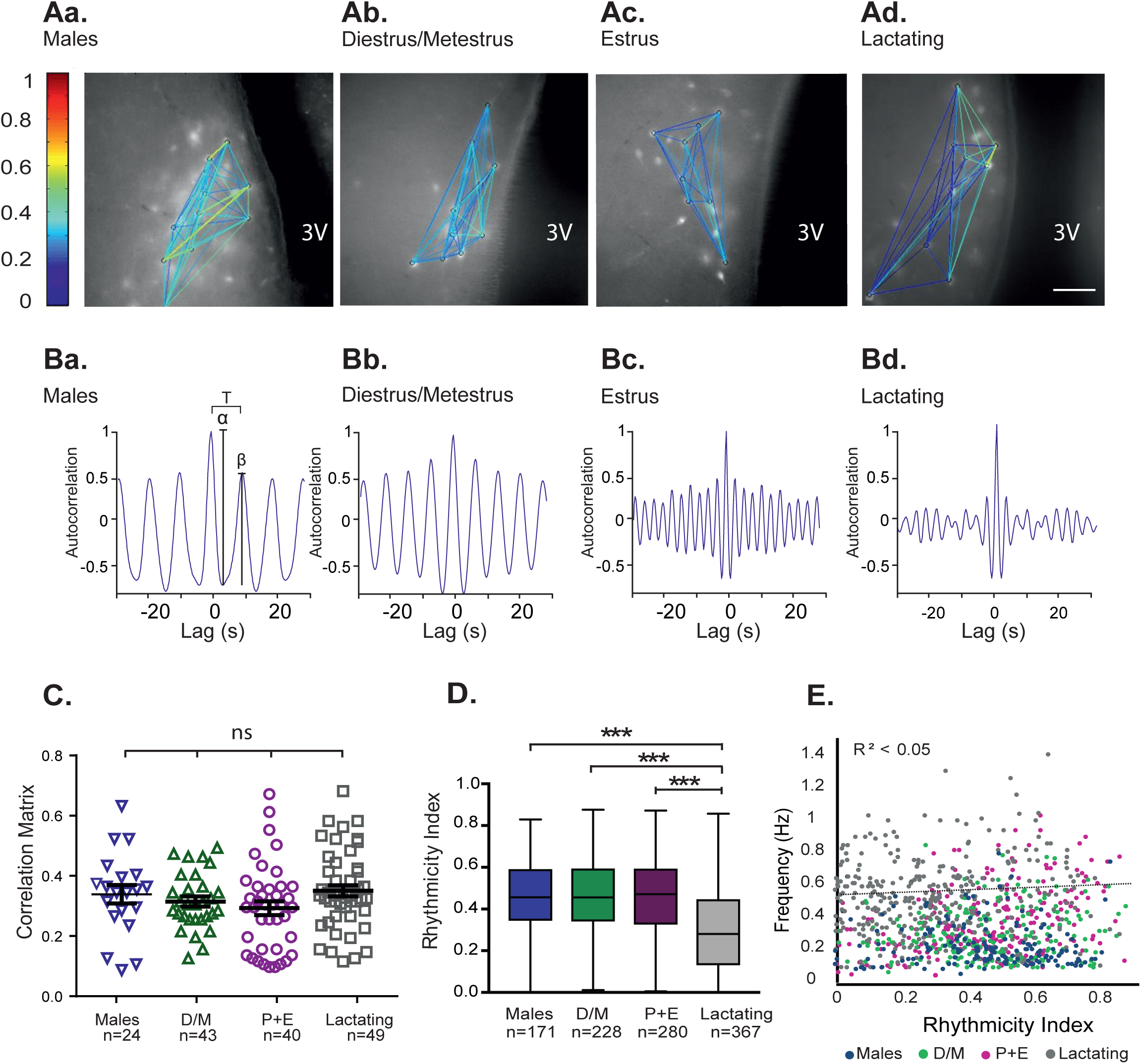
RHYTHMICITY, BUT NOT FUNCTIONAL CONNECTIVITY, OF TIDA NEURONS IS AFFECTED IN LACTATING DAMS. **A.** Representative network maps of the functional connectivity in DAT-GCaMP3 neurons in the dorsomedial arcuate nucleus (dmArc) of adult male (Aa), female in diestrus/metestrus (D/M; Ab), female in estrus (from the proestrus+estrus group; P+E; Ac), and lactating dam (Ad). Degree of correlation between neuron pairs is indicated by colour-coding. **Ba-d.** Representative autocorrelation from one of the neurons from maps shown in *A* for the respective groups. Autocorrelogram in *Ba* illustrates the oscillation period (T), the amplitude of the first trough (α) and the amplitude of the second peak (β); the rhythmicity index (RI) was defined as RI=β/α **C.** There are no significant (ns) differences between the mean values of the correlation matrix between groups. One-way ANOVA with Tukey’s multiple comparison used as *post-hoc* test, “n” given as the number of network maps per hemisphere. **D.** The RI of the lactating group was significantly lower than the RI of the other groups. ***p< 0.001; one-way ANOVA with Tukey’s multiple comparison as *post-hoc* test; “n” given as the number of cells. All other comparisons between groups are non-significant. Scale bar in *Ad* 50 µm for *Aa-d*. 3V, third ventricle. **E.** The correlation between the Ca^2+^ oscillation frequency and the rhythmicity index is low, suggesting that the speed of the oscillation is independent of the rhythmicity.

We also determined whether the oscillation rhythmicity is related to the physiological state and/or to the frequency, by analyzing the auto-correlation of each oscillating cell. Representative auto-correlograms are shown in Figure 3B a-d. During lactation, the rhythmicity index (see *Methods*) of the cells decreased significantly compared to other states, indicating a less regular oscillation pattern (Figure 3D). Notably, rhythmicity did not exhibit a correlation to frequency analyzed across individual cells, suggesting that these factors are independent variables (Figure 3E).

### REVERSIBILITY OF THE TIDA NEURON NETWORK DURING WEANING

Giving birth and nursing are, as a rule, recurring events for mouse dams (and women (Livingston, 2015). At the pituitary level multiple successful pregnancies have been shown to reinforce lactation-associated plastic changes (Hodson et al., 2012). We therefore sought to establish to what extent the frequency changes associated with lactation in TIDA neurons are persistent and/or accentuate with repeated pregnancies. We compared the frequency of Ca^2+^ oscillations in nulliparas, during lactation in primipara mice, in three weeks weaned primipara dams, and in lactating dams following a second pregnancy. Figures 4Aa-d shows representative traces of Ca^2+^ fluctuations in five TIDA neurons from a single slice in a nulliparous (in estrus), a primipara-lactating (same as in Figures 2Ac-d), a weaned female in estrus, and a lactating dam following a second pregnancy (L2). As previously described (*vide supra*) oscillation frequency was significantly higher in the lactating dams. No significant difference was found between the averages for the first and second lactation (Figure 4B). However, there might be subtle differences that do not emerge when the overall mean frequency is compared between slices but are significantly different when compared as individual cells (unpaired, *t*-test; p=0.003). Notably, however, TIDA oscillation frequency in the weaning period was significantly lower than during lactation, and statistically indistinguishable from the estrus period of nulliparous females (Figure 4C). We also compared correlation within the network (Figure 4D) and the rhythmicity (Figure 4E) between the TIDA neurons of nulliparous cycling mice in estrus and of weaned dams in estrus but found no significant difference for either parameter. These data suggest that the lactation-induced resetting of the TIDA network reverses fully once nursing terminates but indicate that there is no “experience” effect of multiple pregnancies at this level of the lactotropic axis.

**FIGURE 4.**
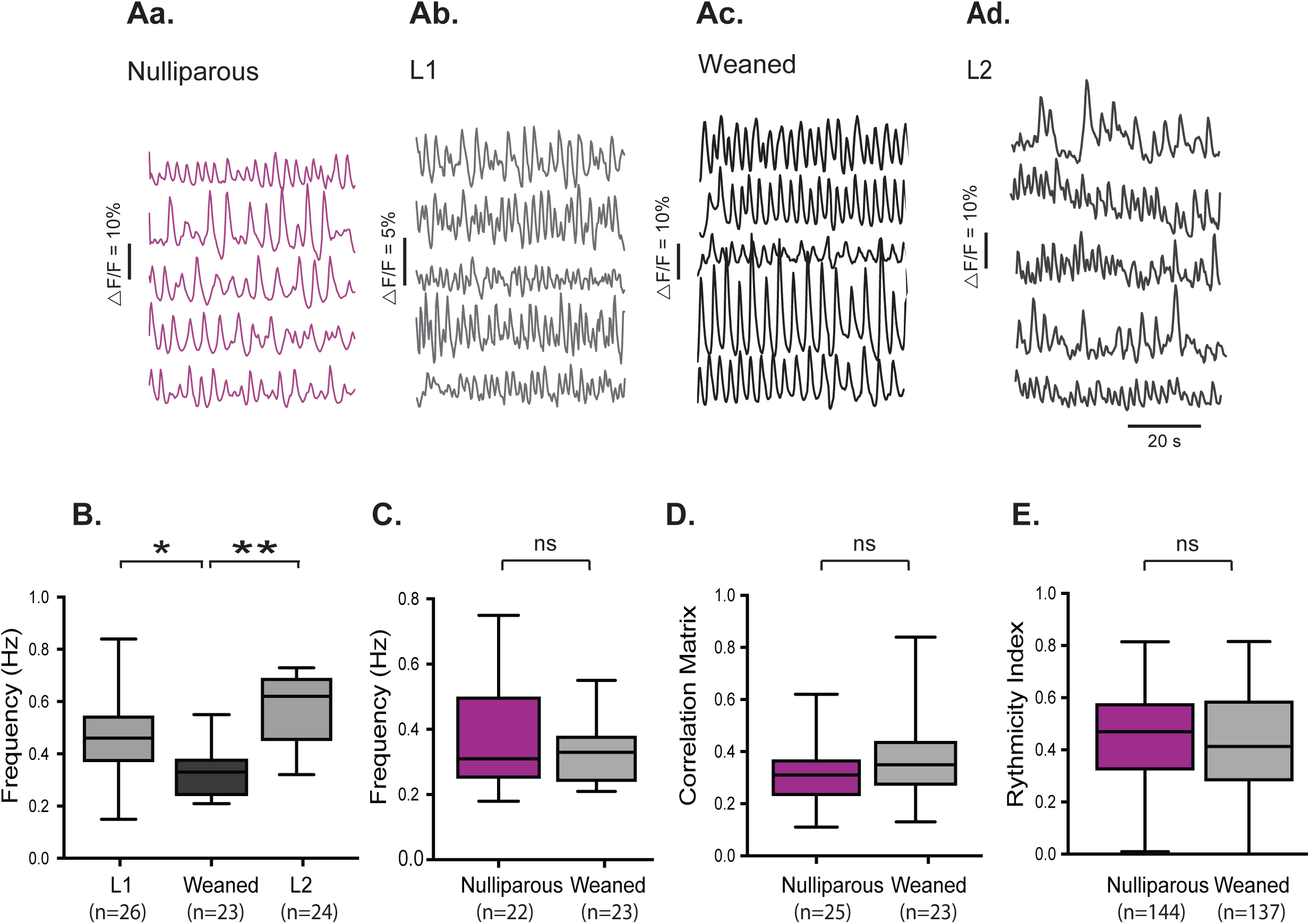
LACTATION-ASSOCIATED NETWORK CHANGES REVERSE UPON WEANING OF PUPS. **A.** Representative concomitant traces of Ca^2+^ fluctuations in DAT-GCaMP3 neurons in the dorsomedial arcuate nucleus (dmArc) from the same slice in nulliparous female in estrus (Aa), primipara lactating (L1) dam (Ab), dam in estrus, three weeks after weaning (Ac), and lactating dam following a second pregnancy (L2; Ad). **B.** Box plot showing average oscillation frequency; note significantly lower frequency in weaned dams compared to L1 and L2. One-way ANOVA with Tukey’ multiple comparison as *post-hoc* test; **p<0.01, *p<0.05. **C.** There is no significant (ns) difference between the average frequency of the Ca^2+^ oscillations between the estrus (nulliparous) group and the weaned females during estrus (p=0.06). **D.** There is no significant difference between the correlation matrix mean between the estrus (nulliparous) group and the weaned females during estrus (p=0.14). “n” given as the number of network maps per hemisphere. **E.** There is no significant difference between the rhythmicity index between the estrus (nulliparous) group and the weaned females during estrus (p=0.30). “n” given as the number of cells. (Aa, Ab same traces as Figures 2Ac, d, respectively.) The number of animals L1: n=5, L2: n=5, Nulliparous: n=7 and Weaned: n=4.

### CHANGES IN INTRINSIC PROPERTIES

Having established that TIDA neurons undergo a reversible acceleration of oscillation frequency during nursing, we next turned our attention to cellular mechanisms that may be involved in this adaptive resetting. This issue was explored by patch clamp whole-cell recordings from TIDA (DAT-GCaMP3 and DAT-tdT) neurons in lactating and cycling nulliparous females. In current clamp recordings, the previously described UP-DOWN state oscillation (Lyons et al., 2010) could clearly be observed in both groups (Figures 5A, B). Notably, the nadir of the oscillating membrane potential of TIDA neurons from nulliparous females was −60.3 ± 1.15 mV (n=11), significantly more hyperpolarized than the TIDA neuron nadir of lactating females (−54.0 ± 1.87 mV, n=10; Figure 5C). To determine if membrane potential was related to oscillation frequency, depolarization was induced by injecting positive constant current into oscillating cells from nulliparous females. This manipulation resulted in faster oscillations (n=11/11, Figure 5A). Contrariwise, hyperpolarization induced by negative current injection to oscillating cells from lactating females resulted in a slower oscillation (n=8/10, Figure 5B). The relationship between imposed (*i.e.* by current injection) nadir potential and oscillation frequency in individual TIDA neurons from nulliparous and lactating females is plotted in Figure 5D. These data suggest that the faster TIDA Ca^2+^ oscillations during lactation may be caused by a depolarizing influence.

**FIGURE 5.**
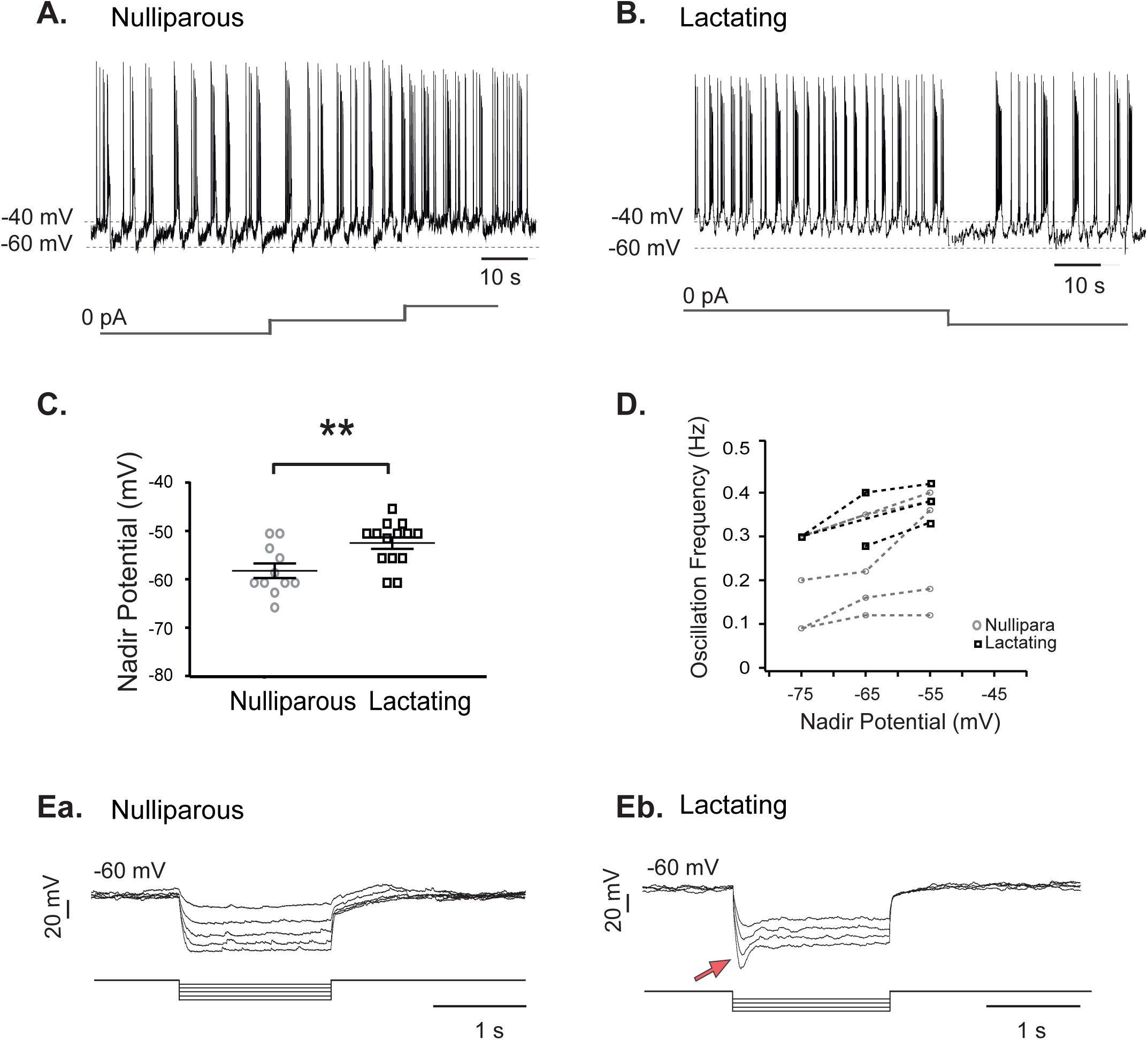
LACTATION-ASSOCIATED INCREASE IN OSCILLATION FREQUENCY IS PARALLELED BY DEPOLARIZATION AND CHANGES IN TIDA MEMBRANE PROPERTIES. **A.** Whole-cell current clamp *in vitro* recording of a DAT-GCaMP3 neuron in the dmArc from a nulliparous female. Membrane potential alternates between hyperpolarized DOWN states and depolarized UP states crowned by action potential discharge. Note increasing frequency of oscillations as depolarizing current of increasing amplitude is injected. **B.** Current clamp recording of a DAT-GCaMP3 neuron from a lactating female. Note faster oscillation frequency that slows down upon injection of hyperpolarizing current. **C.** The mean DOWN state potential (nadir) of DAT-GCaMP3 dmArc neurons from lactating females is more depolarized (−54.00 ± 1.87mV; n=10) than in virgin females (−60.00 ± 1.15mV; n=11). **p<0.01*t*-test. **D.** Nadir potential plotted against oscillation frequency (data from whole-cell recordings in DAT-GCaMP3 neurons). Note linear relationship in both nulliparous and lactating females, and overall higher oscillation frequencies in lactating dams. **Ea.** Electrophysiological recording of a DAT-GCaMP3 neuron from a nulliparous female in the presence of TTX (500 nM), to abolish oscillations and establish a stable baseline, in response to a series of hyperpolarizing square current steps (indicated below). **Eb.** Electrophysiological recording of the voltage response of a DAT-GCaMP3 neuron from lactating female recorded as in Ea. Note presence of depolarizing sag (red arrow), characteristic of I_h_.

Responses of TIDA neurons from nulliparous females to a series of square current injections revealed transient outward rectification, similar to the results reported in rats, where it has been attributed to an A-type K^+^ current (Lyons et al., 2010). In most (n=12/16) cells a post-inhibitory rebound (PIR) was also seen. In no case, however, was a depolarizing sag, indicative of the hyperpolarization-activated non-specific cation current (I_h_), observed (Figures 5Ea). TIDA neurons from lactating females also showed outward rectification (9/16), and PIR was observed in a majority of neurons (n=11/16). Notably, a subpopulation (n=6/16) of TIDA neurons exhibited the depolarizing sag typical of I_h_ (Figures 5Eb). These membrane characteristics showed variable expression among the recorded cells. Thus, the lactation-induced changes in rhythmic activity are paralleled by alterations of specific electrical properties in TIDA neurons, most prominently I_h_, carried by hyperpolarization-activated cyclic nucleotide–gated (HCN) channels.

Expression of HCN channels has been reported in the hypothalamus (Kelly et al., 2013; Loose et al., 1990), but the exact identity of HCN-expressing neurons in this region remains unknown. *In situ* hybridization revealed that mRNA transcripts encoding the HCN subunits1-4 can all be found in TIDA neurons in the arcuate nucleus from nulliparous females during estrus (Figure 6A). Furthermore, the HCN channel auxiliary subunit, the tetratricopeptide repeat-containing Rab8b-interacting protein (TRIP8b; (Lewis et al., 2009; Santoro et al., 2004), was also coexpressed with DAT in the arcuate nucleus (Figure 6A). Notable, a similar expression pattern (while not quantified) was observed in TIDA neurons from lactating dams (Figure 6B). These data indicate that the depolarization sag in TIDA neurons is likely the result of I_h_, but that overt differences in the expression of HCN channels are unlikely to account for the augmentation of this current during lactation.

**FIGURE 6.**
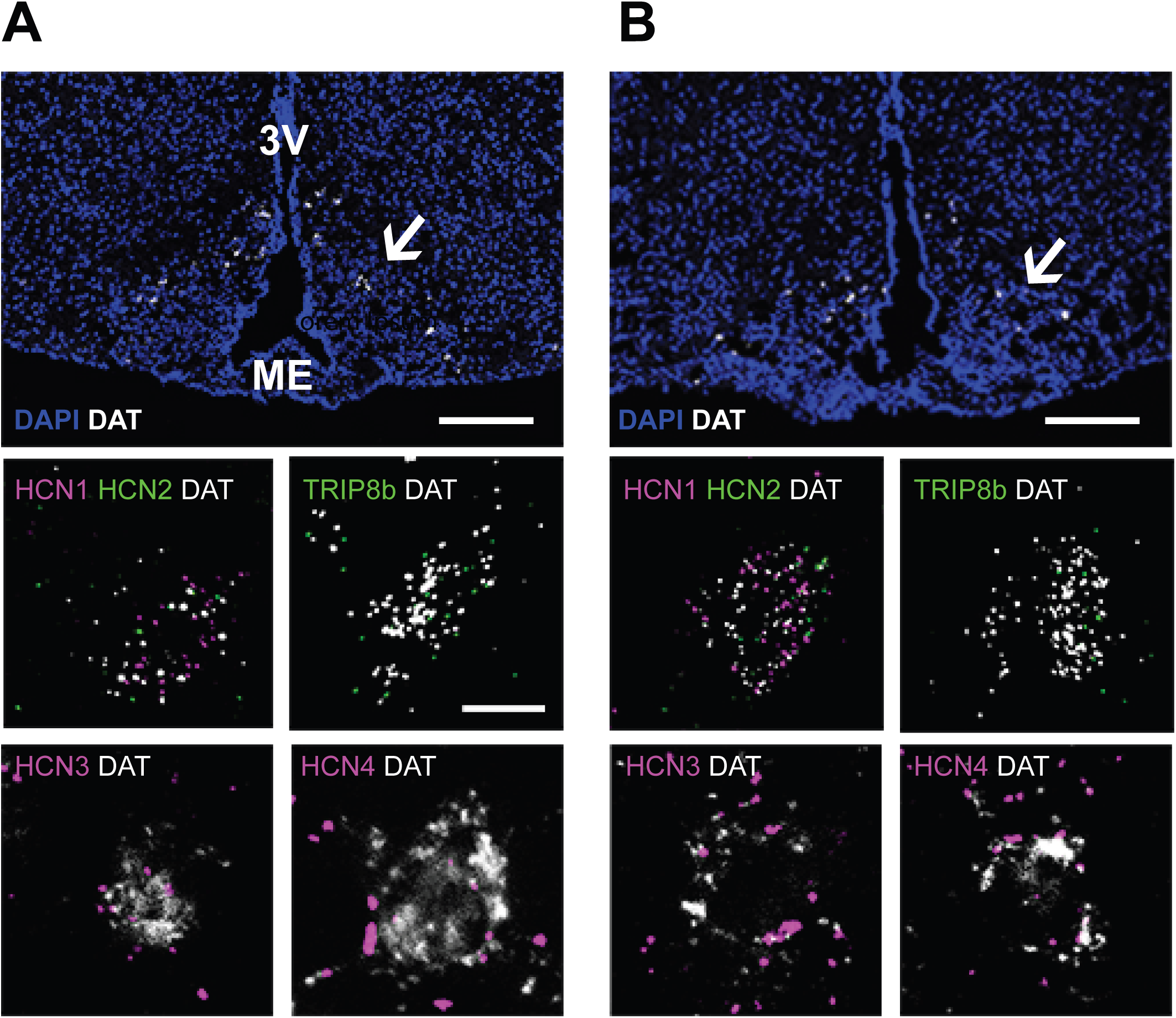
TIDA NEURONS EXPRESS HCN CHANNEL MRNA. **A.** Confocal micrograph from the arcuate nucleus of a female nulliparous mouse in estrus after *in situ* hybridization (RNAScope) performed to detect dopamine transporter (DAT) mRNA (white); section counterstained with DAPI (blue) to visualize anatomy. Note distribution of DAT-expressing TIDA cells in the dorsomedial arcuate nucleus (arrow). Insets show high-magnification examples of TIDA cells co-expressing DAT (white) and (clockwise): the hyperpolarization-activated cyclic nucleotide channel 1 (HCN1; magenta) and HCN2 (green); the HCN channel auxiliary subunit, tetratricopeptide repeat-containing Rab8b-interacting protein (TRIP8b; green); HCN_3_ (magenta); and HCN4 (magenta). **B.** Confocal micrograph organized as in *A*, from the arcuate nucleus of a lactating dam. Note that all RNA transcripts detected in the virgin female TIDA neurons can also be found in the same neurons from a lactating female. Scale bar in A, 100 μ m for all insets. 3V, third ventricle; ME, median eminence.

While I_h_ contributes to rhythmic oscillations in membrane potential in many neurons and other excitable cells (Avella Gonzalez et al., 2015; Bal and McCormick, 1996), earlier studies focusing on male (Lyons et al., 2010) and non-lactating female (Zhang and van den Pol, 2015) rodents have failed to observe current sag in oscillating TIDA cells, similar to our present results in nulliparous mice. The surprising finding of a depolarizing sag in TIDA cells from lactating females (above) prompted us to evaluate the pharmacological identity of this membrane property by applying the selective I_h_ blocker, ZD7288 (Harris and Constanti, 1995). In all six TIDA neurons that displayed a sag upon hyperpolarization, the sag was abolished in the presence of ZD7288 (50 µM; Figure 7Aa), concomitant with a membrane hyperpolarization. Quantification of the change in “sag” amplitude (Δ mV, Figure 7Ab) revealed a significantly greater attenuating effect of ZD7288 in TIDA neurons from lactating dams compared to nulliparous females. Furthermore, the addition of ZD7288 resulted in a more hyperpolarized nadir potential of TIDA cells from lactating dams, but did not affect the nadir potential of TIDA neurons from nulliparous mice (Figure 7Ac). This finding suggests that the depolarized membrane potential of TIDA cells during lactation (Figure 5C, 7Ac) might, at least in part, be related to enhanced I_h_ in this physiological state.

To further validate the presence of I_h_ in TIDA neurons, voltage clamp recordings were performed in lactating dams for comparison to nulliparous females in estrus (Figure 7Ba, Table 1). The response to hyperpolarizing voltage steps, under TTX application, revealed the presence of a ZD7288-sensitive current in both groups (Figure 7Bb). However, the analysis of the currents in response to consecutive command voltage steps indicated a greater sensitivity to ZD7288 in TIDA neurons from lactating mice in comparison with non- lactating mice (Figure 7Bc). Notably, the slope of the I-V curves from TIDA cells during lactation differs significantly from those from nulliparas in control conditions, but the application of ZD7288 eliminates that difference (Figure 7Bd). Thus, although I_h_ can be detected through voltage clamp in nulliparous females, this current is enhanced in lactating dams. The induction or upregulation of this current is thus a candidate to explain, at least in part, the depolarized nadir potential and faster oscillations that TIDA neurons exhibit during nursing.

**Table 1.** Summary of basic electrophysiological properties of TIDA neurons in Nulliparous and Lactating Females. n=5 for each group.

|  | Nulliparous |  | Lactating |  | Significance |
| --- | --- | --- | --- | --- | --- |
| Cell Capacitance | 19,7 ± 2,7 |  | 20,3 ± 3,0 |  | p>0,05 |
|  | Control | ZD | Control | ZD |  |
| Membrane Potential (mV) | -51,4 ± 0,9 | -51,2 ± 0,5 | -42,7 ± 3,3 * | -44,6 ± 2,8 **, ^ | *, **, ^, p<0,05 |
| Input Resistance (GΩ) | 0,7 ± 0,2 | 0,6 ± 0,2 | 2,1 ± 0,5* | 3,5 ± 1,4** ^ | *, **, ^, p<0,05 |
| Experiments were performed under TTX. Data Expressed as Mean ± SE. |  |  |  |  |  |
| *vs. Nulliparous control, **vs. Nulliparous ZD, t-test, ^ vs. LactatingControl, paired-t-test. |  |  |  |  |  |

To test this hypothesis, we assessed if pharmacological blockade of I_h_ affects oscillation frequency recorded at the network level. Thus, ZD7288 (50 µM) was applied during visualization of Ca^2+^ oscillations in DAT-GCaMP3 females. In oscillating TIDA cells from nulliparous females (n=59), the frequency was not significantly affected by I_h_ blockade (Figure 7C). In contrast, in TIDA cells from lactating females (n=72), the mean frequency was significantly reduced in the presence of ZD7288 (Figure 7C), supporting a role for augmented I_h_ in accelerating network rhythms during nursing. Notably, the frequency was significantly different between nulliparous and lactating females under control conditions.

**FIGURE 7.**
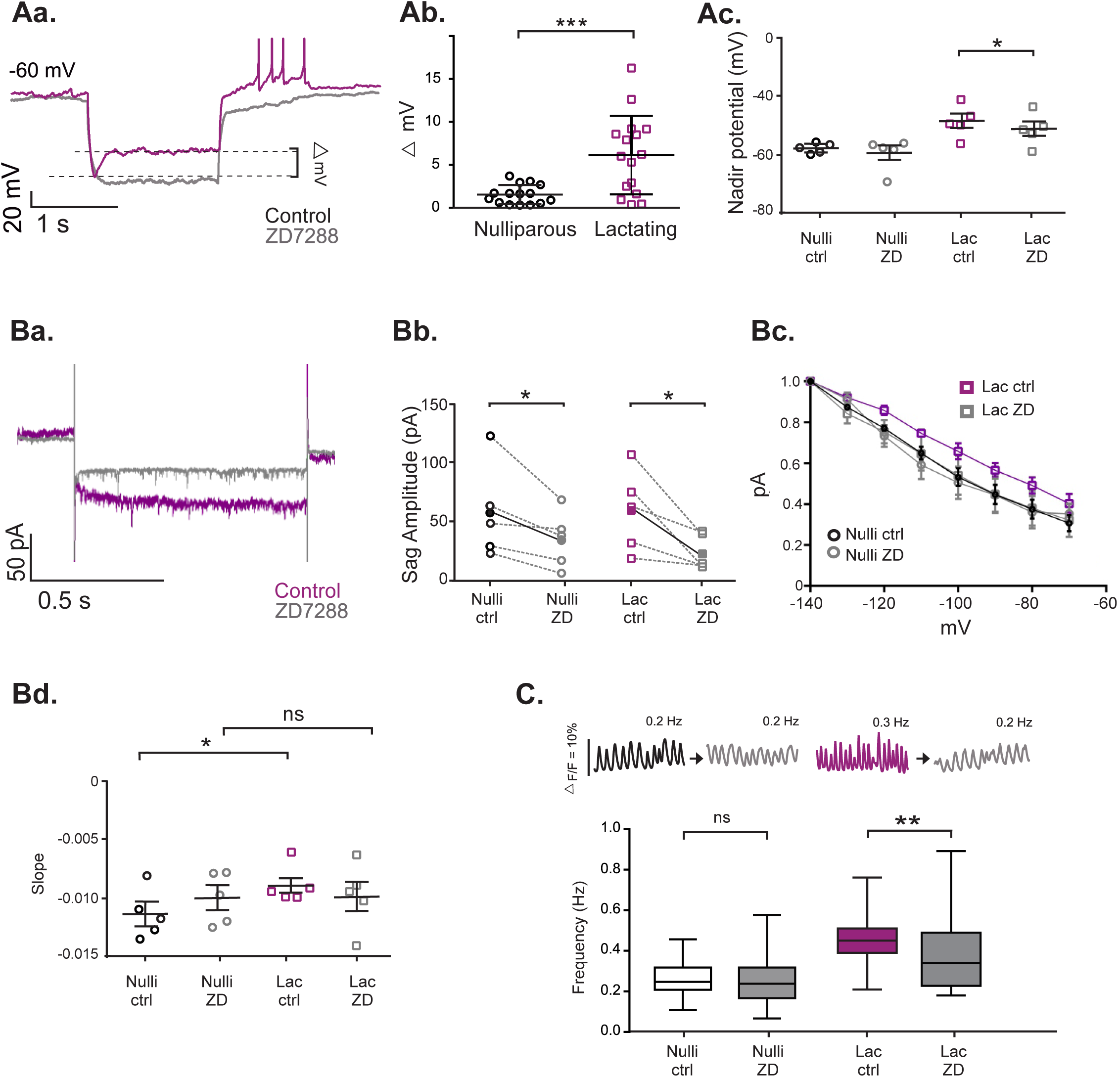
AN INCREASE IN H-CURRENT CONTRIBUTES TO INCREASED TIDA OSCILLATION FREQUENCY DURING LACTATION. **Aa.** Whole-cell current clamp *in vitro* recording of a DAT-GCaMP3 neuron in the dmArc from a lactating dam; negative current step injected to reveal depolarizing sag indicative of h-current (I_h_; magenta trace). Superimposed grey trace shows the same cell recorded in the presence of the I_h_-blocker, ZD7288 (50 µM). Note sag abolished after pharmacological blockade. **Ab.** The voltage difference between the sag and the steady state of a hyperpolarizing step is significantly larger in TIDA neurons from lactating dams compared to nulliparous females. ***P< 0.001, *t*-test. **Ac.** The blockage of I_h_ by ZD7288 decreased the nadir potential in lactating dams (Lac) but not in nulliparous females (Nulli). *p<0.001, paired *t-*test. **Ba.** Representative trace from a whole-cell current clamp recording from a lactating DAT-tdTomato neuron in the dmArc (in the presence of TTX, 500 nM) before (magenta) and after (grey) ZD7288 application (50 µM). **Bb.** ZD7288 decreased the sag amplitude of I_h_ current in both nulliparous and lactating dams. *p<0,05 paired *t-*test. **Bc**. I-V curves from nulliparous and lactating dams in the presence or absence of ZD7288. **Bd**. The comparison of the slope obtained by linear fitting of the I-V curves reveals a difference in lactating dams respect to nulliparous, which is abolished in the presence of Ih blockade by ZD7288, *p<0,05; two-way ANOVA with Tukey’s multiple comparison as *post-hoc* test. **C.** Representative traces of Ca^2+^ fluctuations in a DAT-GCaMP3 neuron before and after ZD7288 application in nulliparous and lactating damn. Box plot illustrating the effect of bath-applied ZD7288 (50 µM) on the frequency of Ca^2+^ oscillations in DAT-GCaMP3 neurons from nulliparous and lactating females. Frequency remains unchanged during pharmacological blockade of I_h_ (paired *t*-test). The frequency was significantly different between control conditions without ZD7288, while in the presence of ZD7288, there is no significant difference between the two groups (one-way ANOVA with Tukey’s multiple comparison as *post-hoc* test).

## DISCUSSION

Few, if any, physiological life events rival the scope and impact of adaptive changes that pregnancy and nursing impose on the body. Virtually all organ systems undergo profound structural and functional modifications to prepare the mother for carrying, giving birth to and nursing her offspring from conception to weaning. In order to organize, coordinate and properly sequence these events, the CNS also undergoes reorganization in key circuits (see (Brunton and Russell, 2008; Elyada and Mizrahi, 2015). In particular, such changes should involve the networks that control the secretion of hormones that define maternal physiology and behaviour. Here, using Ca^2+^ and electrophysiological recordings, we show that TIDA neurons, the main regulator of pituitary prolactin release, shift to faster oscillation frequencies during lactation (but not during the course of the estrous cycle), that this is a reversible phenomenon, and present an upregulation of I_h_ as a candidate mechanism for the resetting of rhythms in the nursing dam.

Earlier work has described altered neurochemistry and modulation of TIDA neurons that allow for a sustained hyperprolactinaemia during lactation that, under most other stages in life, is corrected by powerful feedback mechanisms (see (Le Tissier et al., 2015). Thus, the increase in dopamine output at the neurovascular interface in the median eminence (ME) - that is normally triggered when circulating prolactin rises - is now abolished. Indeed, indices of dopamine release at the ME drop during lactation (Ciofi et al., 1993; Selmanoff and Wise, 1981). This has been attributed to suppressed biosynthesis through down-regulation of expression (Wang et al., 1993) and phosphorylation (Feher et al., 2010; Romano et al., 2013) of tyrosine hydroxylase, and a blunted ability of prolactin to trigger key intracellular signaling pathways (Anderson et al., 2006). Diminished dopamine production is further paralleled by an induction of the peptide, enkephalin (Ciofi et al., 1993; Merchenthaler, 1993; Yip et al., 2019), which may inverse the action of TIDA neurons on prolactin release (see (Le Tissier et al., 2015). We demonstrate here that neuroendocrine adaptation to motherhood also involves changes in the dynamics of TIDA network activity.

Thus, the transition to motherhood appears to involve not only changes in *what* is released from TIDA neurons, and the influence of incoming hormonal signals, but also *how* the neurosecretory signals are released. Interestingly, oscillation frequency was increased only in lactating animals but not in the P+E group, even though the latter period is also characterized by elevated circulating prolactin (Ronnekleiv and Kelly, 1988; Szawka et al., 2007). Thus, it may be speculated that while dynamic recalibration of TIDA rhythms contributes to sustained hyperprolactinaemia during lactation, brief serum peaks of prolactin, such as those within the estrous cycle, result from a qualitatively different modulation of the system. This possibility should, however, be considered with a caveat: if the pre-estrus peak is preceded by a very transient frequency shift that can only be recorded for a brief time, it could have escaped detection in our Ca^2+^ imaging experiments. At the receiving end of the dopamine signal, *i.e.* the pituitary lactotrophs, there is a concomitant consolidation of functional connectivity during lactation (Hodson et al., 2012). A full understanding of how the dopamine-prolactin axis adapts for nursing will thus need to consider the cumulative effects of changes in both sender (TIDA) and receiver (pituitary).

Magnetic resonance imaging has revealed consistent structural modifications in the cerebral cortex of pregnant women (Hoekzema et al., 2017). Overall, published pregnancy- and/or lactation-associated CNS changes generally consist of alterations in structure or gene expression (*e.g.* (Cortes-Sol et al., 2013; Meurisse et al., 2005; Shingo et al., 2003; Theodosis et al., 1986). In comparison, literature describing *functional* puerperal brain plasticity is scarce, with the exception of work identifying how the maternal brain is reorganized to detect and integrate different pup-generated sensory signals (Marlin et al., 2015). These studies offer a precedent for a dynamic reorganization of the dam’s brain, as we report here for a hypothalamic population. In a previous study, where TIDA electrical activity was recorded in loose-patch configuration, oscillation period was not found to be different between male, virgin female and lactating mice (Romano et al., 2013). It is possible that the recording technique, which excludes subthreshold activity, and the smaller number of cells included, precluded detection of the changes we report here. The un-biased nature and large cell yield of Ca^2+^ imaging optimize chances of identifying alterations at the population level.

Our work further identifies augmentation of I_h_ as a candidate mechanism to account for the increased TIDA frequency in dams. This current is implicated in network rhythmicity (see (Luthi and McCormick, 1998), and contributes to depolarization of the resting membrane potential (Maccaferri et al., 1993); see (He et al., 2014), as observed in the lactating mothers (present results). Indeed, we have described a correlation between depolarization and oscillation frequency in mouse TIDA neurons, (Stagkourakis et al., 2018) and the present data show that pharmacological blockade of I_h_ reduces the lactation-associated fast rhythm similar to the rate of nulliparous dams.

The voltage sag typical of I_h_ is not apparent in TIDA neuron current clamp recordings in male juvenile rats (Lyons et al., 2010) or mouse (Zhang and van den Pol, 2015). That does not by necessity mean that the current is fully absent from these neurons. Indeed, indirect evidence supports the functional relevance of I_h_ within the system as application of ZD7288 to rat TIDA cells interferes with the ability of dopamine D2 autoreceptors (D2Rs) to modulate oscillation frequency (Stagkourakis et al., 2016), and in the same system we have observed a ZD7288-induced slowing of network rhythms. The failure to visualize voltage sag in somatic current clamp recordings may reflect a location on dendrites (distant of the location of the recording pipette; (Magee, 1998; Notomi and Shigemoto, 2004; Stuart and Spruston, 1998). In *voltage* clamp recordings, a ZD-sensitive I_h_ component could be seen in nulliparous mouse TIDA neurons (current study). Importantly, however, this component was substantially bigger, and could only be seen in *current* clamp recordings, in TIDA neurons from lactating dams. We thus conclude that I_h_ is augmented during nursing.

The most parsimonious explanation for the appearance of I_h_ in TIDA neurons from lactating dams is an increased transcription of HCN proteins. Indeed, both I_h_ and HCN expression have in other neuronal systems shown to be sensitive to sex steroid status (Piet et al., 2013; Qiao et al., 2013). However, mRNAs for all HCN subunits, as well as the associated HCN protein TRIP8b (Lewis et al., 2009; Santoro et al., 2004), were readily detectable in TIDA neurons in both virgin and lactating females. Though this non-quantitative analysis does not exclude subtle graded changes in transcription, it does suggest that HCN expression is not radically different in virgin and lactating female mice. An alternative scenario thus merits consideration, considering the ultra-short dopaminergic autoinhibitory loop recently described in the TIDA system (Stagkourakis et al., 2016). In the prepubertal male, activation of somatodendritic D2Rs slows TIDA oscillations. Pertinent to the present results, D2Rs can attenuate I_h_ in TIDA neurons (Stagkourakis et al., 2016), respectively, likely through adenylate cyclase inhibition and consequently decreased [cAMP]_intracellular_, a known activator of HCN channels (Parkening et al., 1982; Santoro et al., 1998). Thus, the lactating dam, where TIDA dopamine output has been proposed to be decreased (Demarest et al., 1983; Nagai et al., 1995; Wang et al., 1993), would be likely to experience a decreased D2R-mediated inhibition, leading to increased [cAMP]_intracellular_, which, in turn, would unmask a suppressed I_h_, resulting in depolarization and faster oscillations.

It seems reasonable that at least some of the pregnancy- and lactation-associated brain adaptations need to be reversible to make possible new gravid cycles (up to eight in the female mouse life cycle (Nagai et al., 1995). Yet, examples of such adaptations from the literature have, when assessed, in general demonstrated more or less persistent changes (Cohen et al., 2011; Galindo-Leon et al., 2009; Hoekzema et al., 2017; Macbeth et al., 2008); see (Kinsley et al., 2012)). In contrast, the present results show that after weaning, TIDA oscillations are reset to their pre-pregnancy frequencies. - We also assessed if oscillation frequency differs between a dam’s first and second lactation periods. No significant difference was found when the means of the frequencies between animals were compared. However, when the *t*-test was performed on values from individual cells (*i.e.* a larger sample), the second lactation had was marked by significantly higher oscillation frequencies. This could imply a capacity for “learning” that would improve performance with repeated nursing periods. In humans, reproductive experience impacts on nursing such that mothers produce more milk (Ingram et al., 2001) - although prolactin levels are lower (Dos Santos et al., 2015; Musey et al., 1987) - upon a second and subsequent lactation periods.

In conclusion, the results presented here reveal a new level of plasticity in the neuroendocrine system, where the sustained elevations of prolactin that drive maternal physiology and behaviour in the nursing state are paralleled by a reversible reconfiguration of the hypothalamic circuit that controls its release. Our work identifies a candidate mechanism in the upregulation of I_h_, and shows how the maternal brain resets to prepare for the arrival of offspring.

## ACKNOWLEDGEMENTS

We thank the CLICK facility for access to confocal microscopy equipment and the plasma profiling facility headed by Dr. Jochen Schwenk at Science for Life Laboratory, Stockholm, for the immunoassay tools, and Elin Dahlberg for expert technical assistance. Drs. Nils-Göran Larsson and Ole Kiehn are acknowledged for generously sharing mouse lines, and members of the Broberger laboratory and Dr. Abdel El Manira for helpful discussion. This work was made possible by funding from the European Research Council (ENDOSWITCH 261286), the Swedish Research Council (*Vetenskapsrådet*), the Swedish Brain Foundation (*Hjärnfonden*), the Strategic Research Program for Diabetes Research at Karolinska Institutet, StratNeuro and Novo Nordisk Fonden to CB, and from *Vetenskapsrådet* and *Hjärnfonden* to CTP. JF is supported by a fellowship from the Wenner-Gren Foundations.

## AUTHOR CONTRIBUTIONS

CTP conducted the Ca^2+^ imaging and whole-cell recording experiments, JF conducted and analyzed the whole-cell recording experiments in DAT-tdTomato mice. AH conducted the immunohistochemistry, MJI conducted the serum prolactin analysis. JvL and CTP conducted the MATLAB analysis. CTP and CB designed the experiments and wrote the manuscript. The final version was approved by all authors.

## REFERENCES

Anderson, G.M., Beijer, P., Bang, A.S., Fenwick, M.A., Bunn, S.J., and Grattan, D.R. (2006). Suppression of prolactin-induced signal transducer and activator of transcription 5b signaling and induction of suppressors of cytokine signaling messenger ribonucleic acid in the hypothalamic arcuate nucleus of the rat during late pregnancy and lactation. Endocrinology 147, 4996–5005.

Augustine, R.A., Seymour, A.J., Campbell, R.E., Grattan, D.R., and Brown, C.H. (2018). Integrative neuro-humoral regulation of oxytocin neuron activity in pregnancy and lactation. J Neuroendocrinol.

Avella Gonzalez, O.J., Mansvelder, H.D., van Pelt, J., and van Ooyen, A. (2015). H-Channels Affect Frequency, Power and Amplitude Fluctuations of Neuronal Network Oscillations. Front Comput Neurosci 9, 141.

Bal, T., and McCormick, D.A. (1996). What stops synchronized thalamocortical oscillations? Neuron 17, 297–308.

Bridges, R.S., DiBiase, R., Loundes, D.D., Doherty, P.C., (1985). Prolactin stimulation of maternal behabiour in female rats. Science 227.

Briffaud, V., Williams, P., Courty, J., and Broberger, C. (2015). Excitation of tuberoinfundibular dopamine neurons by oxytocin: crosstalk in the control of lactation. J Neurosci 35, 4229–4237.

Brown, R.S., Kokay, I.C., Phillipps, H.R., Yip, S.H., Gustafson, P., Wyatt, A., Larsen, C.M., Knowles, P., Ladyman, S.R., LeTissier, P., et al. (2016). Conditional Deletion of the Prolactin Receptor Reveals Functional Subpopulations of Dopamine Neurons in the Arcuate Nucleus of the Hypothalamus. J Neurosci 36, 9173–9185.

Brown, R.S.E., Aoki, M., Ladyman, S.R., Phillipps, H.R., Wyatt, A., Boehm, U., and Grattan, D.R. (2017). Prolactin action in the medial preoptic area is necessary for postpartum maternal nursing behavior. Proc Natl Acad Sci U S A 114, 10779–10784.

Brunton, P.J., and Russell, J.A. (2008). The expectant brain: adapting for motherhood. Nat Rev Neurosci 9, 11–25.

Butcher, R.L., Collins, W.E., and Fugo, N.W. (1974). Plasma concentration of LH, FSH, prolactin, progesterone and estradiol-17beta throughout the 4-day estrous cycle of the rat. Endocrinology 94, 1704–1708.

Ciofi, P., Crowley, W.R., Pillez, A., Schmued, L.L., Tramu, G., and Mazzuca, M. (1993). Plasticity in expression of immunoreactivity for neuropeptide Y, enkephalins and neurotensin in the hypothalamic tubero-infundibular dopaminergic system during lactation in mice. J Neuroendocrinol 5, 599–602.

Cohen, L., Rothschild, G., and Mizrahi, A. (2011). Multisensory integration of natural odors and sounds in the auditory cortex. Neuron 72, 357–369.

Cohen-Becker, I.R., Selmanoff, M., Wise, P.M (1986). Inhibitory effects of exogenously induced hyperprolactinemia on the endogenous cyclic release of luteinizing hormone and prolactin in the estyrogen-primed ovarioctomized rat. Endocrinology 119, 1718–1725.

Cortes-Sol, A., Lara-Garcia, M., Alvarado, M., Hudson, R., Berbel, P., and Pacheco, P. (2013). Inner capillary diameter of hypothalamic paraventricular nucleus of female rat increases during lactation. BMC Neurosci 14, 7.

Demarest, K.T., McKay, D.W., Riegle, G.D., and Moore, K.E. (1983). Biochemical indices of tuberoinfundibular dopaminergic neuronal activity during lactation: a lack of response to prolactin. Neuroendocrinology 36, 130–137.

Dorshkind, K., and Horseman, N.D. (2000). The roles of prolactin, growth hormone, insulin-like growth factor-I, and thyroid hormones in lymphocyte development and function: insights from genetic models of hormone and hormone receptor deficiency. Endocr Rev 21, 292–312.

Dos Santos, C.O., Dolzhenko, E., Hodges, E., Smith, A.D., and Hannon, G.J. (2015). An epigenetic memory of pregnancy in the mouse mammary gland. Cell Rep 11, 1102–1109.

Dudley, C.A., Jamison, T.S., and Moss, R.L. (1982). Inhibition of lordosis behavior in the female rat by intraventricular infusion of prolactin and by chronic hyperprolactinemia. Endocrinology 110, 677–679.

Ekstrand, M.I., Terzioglu, M., Galter, D., Zhu, S., Hofstetter, C., Lindqvist, E., Thams, S., Bergstrand, A., Hansson, F.S., Trifunovic, A., et al. (2007). Progressive parkinsonism in mice with respiratory-chain-deficient dopamine neurons. Proc Natl Acad Sci U S A 104, 1325–1330.

Elyada, Y.M., and Mizrahi, A. (2015). Becoming a mother-circuit plasticity underlying maternal behavior. Curr Opin Neurobiol 35, 49–56.

Feher, P., Olah, M., Bodnar, I., Hechtl, D., Bacskay, I., Juhasz, B., Nagy, G.M., and Vecsernyes, M. (2010). Dephosphorylation/inactivation of tyrosine hydroxylase at the median eminence of the hypothalamus is required for suckling-induced prolactin and adrenocorticotrop hormone responses. Brain Res Bull 82, 141–145.

Foo, K.S., Hellysaz, A., and Broberger, C. (2014). Expression and colocalization patterns of calbindin-D28k, calretinin and parvalbumin in the rat hypothalamic arcuate nucleus. J Chem Neuroanat 61-62, 20–32.

Fox, S.R., Hoefer, M.T., Bartke, A., Smith, M.S., (1987). Suppresion of pulsatile LH secretion, pituitary GhRH receptor content and pituitary responsiveness to GnRG by hyperprolactenemia in the male rat. Neuroendocrinology 46, 350–359.

Freeman, M.E., Kanyicska, B., Lerant, A., and Nagy, G. (2000). Prolactin: structure, function, and regulation of secretion. Physiol Rev 80, 1523–1631.

Galindo-Leon, E.E., Lin, F.G., and Liu, R.C. (2009). Inhibitory plasticity in a lateral band improves cortical detection of natural vocalizations. Neuron 62, 705–716.

Grattan, D.R. (2015). 60 YEARS OF NEUROENDOCRINOLOGY: The hypothalamo-prolactin axis. J Endocrinol 226, T101–122.

Grattan, D.R., Steyn, F.J., Kokay, I.C., Anderson, G.M., and Bunn, S.J. (2008). Pregnancy-induced adaptation in the neuroendocrine control of prolactin secretion. J Neuroendocrinol 20, 497–507.

Guillou, A., Romano, N., Steyn, F., Abitbol, K., Le Tissier, P., Bonnefont, X., Chen, C., Mollard, P., and Martin, A.O. (2015). Assessment of lactotroph axis functionality in mice: longitudinal monitoring of PRL secretion by ultrasensitive-ELISA. Endocrinology 156, 1924–1930.

Harris, N.C., and Constanti, A. (1995). Mechanism of block by ZD 7288 of the hyperpolarization-activated inward rectifying current in guinea pig substantia nigra neurons in vitro. J Neurophysiol 74, 2366–2378.

He, C., Chen, F., Li, B., and Hu, Z. (2014). Neurophysiology of HCN channels: from cellular functions to multiple regulations. Prog Neurobiol 112, 1–23.

Hodson, D.J., Schaeffer, M., Romano, N., Fontanaud, P., Lafont, C., Birkenstock, J., Molino, F., Christian, H., Lockey, J., Carmignac, D., et al. (2012). Existence of long-lasting experience-dependent plasticity in endocrine cell networks. Nat Commun 3, 605.

Hoekzema, E., Barba-Muller, E., Pozzobon, C., Picado, M., Lucco, F., Garcia-Garcia, D., Soliva, J.C., Tobena, A., Desco, M., Crone, E.A., et al. (2017). Pregnancy leads to long-lasting changes in human brain structure. Nat Neurosci 20, 287–296.

Ingram, J., Woolridge, M., and Greenwood, R. (2001). Breastfeeding: it is worth trying with the second baby. Lancet 358, 986–987.

Kalyani, M., Hasselfeld, K., Janik, J.M., Callahan, P., and Shi, H. (2016). Effects of High-Fat Diet on Stress Response in Male and Female Wildtype and Prolactin Knockout Mice. PLoS One 11, e0166416.

Katz, B., and Miledi, R. (1970). Further study of the role of calcium in synaptic transmission. J Physiol 207, 789–801.

Kelly, M.J., Zhang, C., Qiu, J., and Ronnekleiv, O.K. (2013). Pacemaking kisspeptin neurons. Exp Physiol 98, 1535–1543.

Kinsley, C.H., Franssen, R.A., and Meyer, E.A. (2012). Reproductive experience may positively adjust the trajectory of senescence. Curr Top Behav Neurosci 10, 317–345.

Le Tissier, P.R., Hodson, D.J., Martin, A.O., Romano, N., and Mollard, P. (2015). Plasticity of the prolactin (PRL) axis: mechanisms underlying regulation of output in female mice. Adv Exp Med Biol 846, 139–162.

Lerant, A., and Freeman, M.E. (1998). Ovarian steroids differentially regulate the expression of PRL-R in neuroendocrine dopaminergic neuron populations: a double label confocal microscopic study. Brain Res 802, 141–154.

Levine, J.D., Funes, P., Dowse, H.B., and Hall, J.C. (2002). Resetting the circadian clock by social experience in Drosophila melanogaster. Science 298, 2010–2012.

Lewis, A.S., Schwartz, E., Chan, C.S., Noam, Y., Shin, M., Wadman, W.J., Surmeier, D.J., Baram, T.Z., Macdonald, R.L., and Chetkovich, D.M. (2009). Alternatively spliced isoforms of TRIP8b differentially control h channel trafficking and function. J Neurosci 29, 6250–6265.

Livingston, G. (2015). Family Size Among Mothers. Pew Research Center.

Loose, M.D., Ronnekleiv, O.K., and Kelly, M.J. (1990). Membrane properties and response to opioids of identified dopamine neurons in the guinea pig hypothalamus. J Neurosci 10, 3627–3634.

Lorang, D., Amara, S.G., and Simerly, R.B. (1994). Cell-type-specific expression of catecholamine transporters in the rat brain. J Neurosci 14, 4903–4914.

Luthi, A., and McCormick, D.A. (1998). H-current: properties of a neuronal and network pacemaker. Neuron 21, 9–12.

Lyons, D.J., Ammari, R., Hellysaz, A., and Broberger, C. (2016). Serotonin and Antidepressant SSRIs Inhibit Rat Neuroendocrine Dopamine Neurons: Parallel Actions in the Lactotrophic Axis. J Neurosci 36, 7392–7406.

Lyons, D.J., and Broberger, C. (2014). TIDAL WAVES: Network mechanisms in the neuroendocrine control of prolactin release. Front Neuroendocrinol 35, 420–438.

Lyons, D.J., Horjales-Araujo, E., and Broberger, C. (2010). Synchronized network oscillations in rat tuberoinfundibular dopamine neurons: switch to tonic discharge by thyrotropin-releasing hormone. Neuron 65, 217–229.

Macbeth, A.H., Gautreaux, C., and Luine, V.N. (2008). Pregnant rats show enhanced spatial memory, decreased anxiety, and altered levels of monoaminergic neurotransmitters. Brain Res 1241, 136–147.

Maccaferri, G., Mangoni, M., Lazzari, A., and DiFrancesco, D. (1993). Properties of the hyperpolarization-activated current in rat hippocampal CA1 pyramidal cells. J Neurophysiol 69, 2129–2136.

Magee, J.C. (1998). Dendritic hyperpolarization-activated currents modify the integrative properties of hippocampal CA1 pyramidal neurons. J Neurosci 18, 7613–7624.

Marlin, B.J., Mitre, M., D’Amour J, A., Chao, M.V., and Froemke, R.C. (2015). Oxytocin enables maternal behaviour by balancing cortical inhibition. Nature 520, 499–504.

McLean, A.C., Valenzuela, N., Fai, S., and Bennett, S.A. (2012). Performing vaginal lavage, crystal violet staining, and vaginal cytological evaluation for mouse estrous cycle staging identification. J Vis Exp, e4389.

Merchenthaler, I. (1993). Induction of enkephalin in tuberoinfundibular dopaminergic neurons during lactation. Endocrinology 133, 2645–2651.

Meurisse, M., Gonzalez, A., Delsol, G., Caba, M., Levy, F., and Poindron, P. (2005). Estradiol receptor-alpha expression in hypothalamic and limbic regions of ewes is influenced by physiological state and maternal experience. Horm Behav 48, 34–43.

Morishige, W.K., and Rothchild, I. (1974). Temporal aspects of the regulation of corpus luteum function by luteinizing hormone, prolactin and placental luteotrophin during the first half of pregnancy in the rat. Endocrinology 95, 260–274.

Musey, V.C., Collins, D.C., Brogan, D.R., Santos, V.R., Musey, P.I., Martino-Saltzman, D., and Preedy, J.R. (1987). Long term effects of a first pregnancy on the hormonal environment: estrogens and androgens. J Clin Endocrinol Metab 64, 111–118.

Nagai, J., Lin, C.Y., and Sabour, M.P. (1995). Lines of mice selected for reproductive longevity. Growth Dev Aging 59, 79–91.

Notomi, T., and Shigemoto, R. (2004). Immunohistochemical localization of Ih channel subunits, HCN1-4, in the rat brain. J Comp Neurol 471, 241–276.

Parkening, T.A., Collins, T.J., and Smith, E.R. (1982). Plasma and pituitary concentrations of LH, FSH, and prolactin in aging C57BL/6 mice at various times of the estrous cycle. Neurobiol Aging 3, 31–35.

Piet, R., Boehm, U., and Herbison, A.E. (2013). Estrous cycle plasticity in the hyperpolarization-activated current ih is mediated by circulating 17beta-estradiol in preoptic area kisspeptin neurons. J Neurosci 33, 10828–10839.

Qiao, G.F., Qian, Z., Sun, H.L., Xu, W.X., Yan, Z.Y., Liu, Y., Zhou, J.Y., Zhang, H.C., Wang, L.J., Pan, X.D., et al. (2013). Remodeling of hyperpolarization-activated current, Ih, in Ah-type visceral ganglion neurons following ovariectomy in adult rats. PLoS One 8, e71184.

Qundos, U., Johannesson, H., Fredolini, C., O’Hurley, G., Branca, R., Uhlén, M., Wiklund, F., Bjartell, A., Nilsson, P., and Schwenk, J.M. (2014). Analysis of plasma from prostate cancer patients links decreased carnosine dipeptidase 1 levels to lymph node metastasis. Translational Proteomics 2, 14–24.

Romano, N., Yip, S.H., Hodson, D.J., Guillou, A., Parnaudeau, S., Kirk, S., Tronche, F., Bonnefont, X., Le Tissier, P., Bunn, S.J., et al. (2013). Plasticity of hypothalamic dopamine neurons during lactation results in dissociation of electrical activity and release. J Neurosci 33, 4424–4433.

Ronnekleiv, O.K., and Kelly, M.J. (1988). Plasma prolactin and luteinizing hormone profiles during the estrous cycle of the female rat: effects of surgically induced persistent estrus. Neuroendocrinology 47, 133–141.

Sairenji, T.J., Ikezawa, J., Kaneko, R., Masuda, S., Uchida, K., Takanashi, Y., Masuda, H., Sairenji, T., Amano, I., Takatsuru, Y., et al. (2017). Maternal prolactin during late pregnancy is important in generating nurturing behavior in the offspring. Proc Natl Acad Sci U S A 114, 13042–13047.

Santoro, B., Liu, D.T., Yao, H., Bartsch, D., Kandel, E.R., Siegelbaum, S.A., and Tibbs, G.R. (1998). Identification of a gene encoding a hyperpolarization-activated pacemaker channel of brain. Cell 93, 717–729.

Santoro, B., Wainger, B.J., and Siegelbaum, S.A. (2004). Regulation of HCN channel surface expression by a novel C-terminal protein-protein interaction. J Neurosci 24, 10750–10762.

Sauve, D., Woodside, B., (1996). The effect of central administration of prolactin on food intake in virgin female ratos is dose-dependent, occurs in the absence of ovarian hormones and the latency to onset varies with feeding regimen. Brain Res 729, 75–81.

Selmanoff, M., and Wise, P.M. (1981). Decreased dopamine turnover in the median eminence in response to suckling in the lactating rat. Brain Res 212, 101–115.

Shingo, T., Gregg, C., Enwere, E., Fujikawa, H., Hassam, R., Geary, C., Cross, J.C., and Weiss, S. (2003). Pregnancy-stimulated neurogenesis in the adult female forebrain mediated by prolactin. Science 299, 117–120.

Smedler, E., Malmersjo, S., and Uhlen, P. (2014). Network analysis of time-lapse microscopy recordings. Front Neural Circuits 8, 111.

Stagkourakis, S., Kim, H., Lyons, D.J., and Broberger, C. (2016). Dopamine Autoreceptor Regulation of a Hypothalamic Dopaminergic Network. Cell Rep.

Stagkourakis, S., Perez, C.T., Hellysaz, A., Ammari, R., and Broberger, C. (2018). Network oscillation rules imposed by species-specific electrical coupling. Elife 7.

Stricker, P.G., F., (1928). Action du lobe anterieur de l’hypophyse sur la montee laiteuse. Compt Rend Soc Biol 99, 1978–1980.

Stuart, G., and Spruston, N. (1998). Determinants of voltage attenuation in neocortical pyramidal neuron dendrites. J Neurosci 18, 3501–3510.

Szawka, R.E., Rodovalho, G.V., Helena, C.V., Franci, C.R., and Anselmo-Franci, J.A. (2007). Prolactin secretory surge during estrus coincides with increased dopamine activity in the hypothalamus and preoptic area and is not altered by ovariectomy on proestrus. Brain Res Bull 73, 127–134.

Theodosis, D.T., Chapman, D.B., Montagnese, C., Poulain, D.A., and Morris, J.F. (1986). Structural plasticity in the hypothalamic supraoptic nucleus at lactation affects oxytocin-, but not vasopressin-secreting neurones. Neuroscience 17, 661–678.

Wang, H.J., Hoffman, G.E., and Smith, M.S. (1993). Suppressed tyrosine hydroxylase gene expression in the tuberoinfundibular dopaminergic system during lactation. Endocrinology 133, 1657–1663.

Yip, S.H., Romano, N., Gustafson, P., Hodson, D.J., Williams, E.J., Kokay, I.C., Martin, A.O., Mollard, P., Grattan, D.R., and Bunn, S.J. (2019). Elevated Prolactin during Pregnancy Drives a Phenotypic Switch in Mouse Hypothalamic Dopaminergic Neurons. Cell Rep 26, 1787–1799 e1785.

Zhang, X., and van den Pol, A.N. (2015). Dopamine/Tyrosine Hydroxylase Neurons of the Hypothalamic Arcuate Nucleus Release GABA, Communicate with Dopaminergic and Other Arcuate Neurons, and Respond to Dynorphin, Met-Enkephalin, and Oxytocin. J Neurosci 35, 14966–14982.

